# Time and spatial trends in landing per unit of effort as support to fisheries management in a multi-gear coastal fishery

**DOI:** 10.1101/2021.10.04.463092

**Authors:** P. Leitão, L. Sousa, M. Castro, A. Campos

## Abstract

Landings by the multi-gear coastal fleet operating off the Portuguese continental coast include near 300 species, from which only a few are the object of management plans. In this study, daily landings (kg trip^_1^) are used, along with an effort indicator, vessel length overall (LoA), to obtain landings per unit of effort (LPUE) as a proxy for the species relative abundance for a total of 48 species. LPUE indices were then used as a response variable in linear models where year (2012-2016), season, region (north and south) and NAO index were included as explanatory variables. Season and region effects were found to significantly affect species abundance for a total of 41 and 40 species respectively, while interannual effects were found to be significant for 19 species, and finally, the NAO index for 3 species. Global LPUE density maps are presented for a number of selected species and a subsample of trips where VMS records were available. For the species analysed, it is proposed that geographic and seasonal changes in LPUE indexes can be used to understand trends in abundance and obtain information that can be used in support of the definition of regional management plans.

## INTRODUCTION

In the European Union (EU), the principles underlying the Common Fisheries Policy (CFP) focus on long-term sustainability of marine resources. Fisheries were recognized to be dependent on healthy marine ecosystems, requiring the integration of the fisheries sector in other policies dealing with marine activities, namely the Integrated Maritime Policy (IMP) and its environmental pillar, the Marine Strategy Framework Directive (MSFD). An important aspect of these policies is the implementation of the ecosystem-based approach to fisheries management (EAFM), which became a central issue under the CFP, requiring information on the state of the marine environment, including indicators of fishing pressure, both for target and non-target species. Emphasis was placed on the need for a regionalized approach to fisheries management, with the establishment of fishery-based management plans and mitigation measures to be tailored to specific fisheries. Single species management will be replaced by multi-species long-term plans, while the spatial component plays an increasingly important role as a broader set of ecosystem interactions need to be considered.

For a sustainable fishing, management measures have evolved from control of catches (e.g., total allowable captures (TACs) and quotas) to fishing effort limitations and technical conservation measures, including gear restrictions, species minimum conservation reference sizes (MCRS) and real time closures (1). A combination of these types of measures is used in European fishery policies. The EAFM calls upon a fleet- and area-based approach to fisheries management (Council Regulation (EC) No 1343/2007) increasingly requiring the integration of fisheries-dependent data.

The evolution of methodologies for the assessment and management of fishing resources took place in parallel with important changes in the collection of data. Currently, most of the information relevant to fisheries management in the EU has been obtained either through vessel research surveys or fleet monitoring programs at the scope of the Data Collection Framework (DCF). In the last two decades, new mandatory procedures evolved, to improve and standardize fisheries data throughout the EU (2). Large and more reliable datasets are generated, and statistical modelling can then be applied to retrieve crucial fisheries information such as the identification of exploited species and their distribution (3–5). The International Council for the Exploration of the Sea (ICES), the entity responsible for providing advice on fisheries issues to the EU, has identified as a major objective, in its Science Plan, the development of effective mechanisms to use monitoring and surveillance data to support scientific advice (6).

The use of fisheries-dependent data is particularly useful in fisheries advice in fisheries where a high number of commercial species are exploited, from which only some are subject to formal assessment by ICES, resulting in TACs and quotas. For these areas, alternative management methods should be developed including area-specific management based on easily accessible fisheries dependent data. This approach assumes particular importance for fleets not covered by onboard sampling programs (7,8).

Traditional stock assessment has relied on fisheries independent surveys to evaluate the biomass of exploited resources (9,10). In the absence of this information, fisheries dependent data, such as catch per unit of effort (CPUE), represents the best available indicator of abundance (11–13). CPUE data are mainly gathered through national sampling programs where onboard observers register, for specific trips, the catch composition (number, weight and sizes of target and bycatch species) and respective fishing effort (hours of fishing, type, number, and size of gears).

For data deficient fleets, landings, instead of catches, can be used to obtain an alternative abundance index, landings per unit of effort (LPUE) (14,15). Landings, combined with georeferenced data on the fishing activity, can provide information on stock trends at a regional level, allowing closures in space and time (16). Near real-time closures (RTCs) based on daily high-definition fishing maps of CPUE are beneficial to fisheries, in particular for heavily fished demersal species (17,18). Iceland and Scotland rely on georeferenced catch data to define RTCs, limiting fishing in particular areas and ensuring sustainability (17).

The multi-gear fishing fleet operating on the Portuguese continental coast (ICES Division IXa) accounts for 96% of the total number of vessels, employing 69% of fishers and being responsible for 31% of the landings in weight and 59% in value (19). This complex fleet operates year-round, over a great variety of ecosystems, adapted to regional and seasonal availability of resources. Most vessels are licensed for multiple fixed fishing gears, such as gillnets and trammel nets, longlines, traps and pots, to capture a great diversity of benthic, demersal and pelagic species (fish, shellfish, cephalopods, and crustaceans). Due to limitations of crew size and well-being conditions, only a small fringe of the multi-gear fleet can accommodate onboard observers and is monitored through the EU Data Collection Framework (DCF) program (20).

The objective of this study is to illustrate the utility of using daily landing records and a measure of effort to obtain LPUE indexes as a proxy of abundance for a high number of species captured by a multi-gear, multi-species fleet, and relate LPUE with environmental information. Short term trends (2012-2016) were obtained for a large number of species. For a selection of species, the utility of using available georeferenced information given by VMS data was demonstrated in mapping LPUE, illustrating spatial and time patterns in exploitation that can contribute do define regional management plans.

## MATERIALS AND METHODS

### Data sources

The Portuguese multi-gear fleet, operating mainly nets, traps and longlines, account for low bycatch when compared to purse seine and bottom trawlers, and generally take an opportunistic “all species are valuable” approach, discarding only species interdicted or those with no commercial value (21–23). In such conditions, landings and catches are closely related and, in the absence of catch per unit of effort (CPUE), landings per unit of effort (LPUE) are considered as a good proxy for species abundance (24).

The data analysis refers to the multi-gear coastal fleet operating off the continental Portuguese coast, comprising a total of 492 vessels greater than 9 meters in length. Although most vessels use different types of static gear, a small fleet sub-segment uses dredges all year round to exclusively target bivalves. For each fishing trip, the data consisted of daily landings by species and technical characteristics of the vessels. For a sample of 165 vessels greater than 15 meters, VMS records associated with the fishing trips were also available.

#### Landings data

Daily landings included date, landing port, vessel ID, species identification (common name and the 3-alpha FAO code) and landings in weight (kg) and value (€ kg^-1^). for the 492 vessels belonging to the multi-gear coastal fleet. The original landings dataset comprised a total of 257,134 fishing trips and 297 species. Three criteria were applied in sequence, to eliminate very rare species and/or trips with very low catches: (1) selection of the 100 species with higher landing frequencies, (2) based on the previous 100 species, selection of the 50 species with higher total landings (in weight), (3) for each of these 50 species, selection of the trips that landed more than 10% of the average daily landing of that species. In addition, the genus *Microchirus* was removed due to duplicated records as genus and species. The application of these criteria resulted in a total of 48 species captured in 247,252 trips, representing a loss of 3.8% of the original number of trips.

#### Vessel characteristics and VMS data

The technical characteristics of the vessels comprised length overall (LoA), gross tonnage (GT), vessel power in kW, year of construction and port of register.

Regarding VMS, the coastal fleet of interest is composed of vessels with length overall (LoA) greater than 9 meters, but VMS systems are only installed in some vessels larger than 12 meters and in all vessels larger than 15 meters. Vessels between 12 and 15 meters are exempt from the obligation to have the monitoring equipment if they operate exclusively within territorial waters or spend less than 24 hours at sea. Due to this, of the 492 vessels in the multi-gear coastal fleet only, 165 were equipped with VMS, representing approximately 90 000 trips.

VMS positioning data for each vessel consists of a succession of geographical locations (latitude, longitude), timestamp, speed, and course, received by a ‘blue-box’ (satellite-tracking device installed on board the fishing vessels). This information is transmitted via satellite to the Fisheries Control Centre every 2 hours.

At an initial data processing stage, VMS data were filtered to exclude records with duplicate or erroneous values. The data analysis proceeded with the identification of fishing trips (FT) for the 165 vessels. This identification was carried out by partitioning the VMS data into sections starting with departure from a port and ending at the arrival to the same or a different port. Each one of these sections, corresponding to a fishing trip, is associated to a landing declaration. The objective was, for each species, to provide spatial-temporal information on fishing grounds and relative abundance, together with information on fishing effort.

The VMS data, landings, and vessel technical characteristics were provided in an anonymized format (each vessel was attributed a code) by the Directorate-General for Natural Resources, Safety and Maritime Services (DGRM).

#### Geographical, temporal, and environmental variables

Previous studies on spatial distribution of fish assemblages off the Portuguese continental shelf, based on the analysis of trawl surveys (25), demonstrated the existence of two main biological regions, separated by a boundary located around the Nazaré Canyon, a steep-side valley that represents not only a physical obstacle for fish communities but also separates areas that are geologically and environmentally different. The southern continental shelf is narrow and affected by weak outward winds, while the northern shelf is wider and is influenced by southward coast winds, creating the conditions for upwelling and higher primary production and leading to higher pelagic fish abundance (26,27). This boundary was considered in the present study, using the landing port to associate the fishing activity to a region, north or south, in relation to the Nazaré canyon.

Two variables related with time were considered as possibly influencing LPUE, year (2012 to 2016) and season (winter from January to March (1 – 3), spring from April to June (4 – 6); summer from July to September (7 – 9) and fall from October to December (10 – 12)).

An environmental variable was included, the North Atlantic Oscillation (NAO) index. This index is related to the difference between low atmospheric pressure at high latitudes and high atmospheric pressure at low latitudes in the North Atlantic and its magnitude depends on the choice of the sampling sites where high and low atmospheric pressures are measured, as well as on the seasons considered (28). In this work we used the NAO index proposed by Hurrell (1995) that considers the difference of normalized sea level pressure between the Azores (high) and Iceland (low). The NAO influences the direction and strength of western winds, and high index values during the previous winter may have a positive effect on primary production (30,31), positively affecting zooplankton abundance and favouring larval survival and fish recruitment (32,33). A time-lag (δ in years) was considered for each species, calculated as the number of years between larval phase and recruitment to fisheries; the LPUE of a given year Y was thus associated with the NAO index of year Y-δ (table with lag values in supplementary material).

### Standardized fishing effort and LPUE

Detailed information on fishing effort was not available for the fleet of interest. However, Portuguese fishing regulations, Ordinance nº 1102-H/2000, defines, for vessels larger than 9 meters in length overall (LoA), six vessel size categories, with corresponding maximum numbers of gears, where vessels with higher length overall (LoA) are allowed 2 to 3 times higher fishing effort due to longer fleets and more traps. Thus, vessel length was considered a proxy for fishing capacity. A second indicator is the engine power, commonly used to standardize fishing effort in trawlers (20). These two potential indicators of the vessel’s fishing capacity (length overall and engine power) were investigated through their correlation with landed weight. A stronger relationship was present between LoA and landed weight (deviance explained was 6.24% for LoA and 0.13% for horsepower).

Daily landings per unit of effort (LPUE_*s,d*_) for each species was calculated through equation 1:

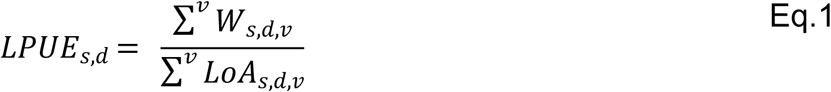

where the landed weight (*W; kilograms*) for species *s*, day *d* and vessel *v*, is summed for all vessels for which landings for those species were equal or higher than 10% of the total landings of that vessel and divided by the sum of the vessel LoA (meters).

### Statistical methods

General linear models were used in this study to evaluate the influence of multiple variables on LPUE. Those relationships have been used in this context to establish causal relationships and predicting future outcomes (34).

The variables used in the statistical analysis were: LPUE per day and species as response variable; year and NAO index as continuous explanatory variables; and region and season as categorical explanatory variables.

The log_10_(LPUE) was the response variable for the model:

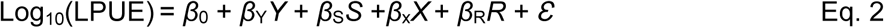

where *Y* the YEAR, *S* is SEASON, *X* the NAO INDEX, *R* the REGION and *ε* the associated error, and LPUE for each species is in kg per meter boat length per day.

The model was adjusted for each of the 48 species selected. The p-values of the specific terms of the model, with a Bonferroni correction for multiple tests, were used to assess statistical significance and strength of the explanatory variables (35). Significance was discussed only for more important variables with p-values less than 0.01.

Database setup and statistical analysis were carried out using R version 3.6.3 on RStudio.

### Mapping LPUE

Maps of fishing abundance were obtained for a number of selected species, using QGIS version 3.10. Daily landings of a particular species and vessel were assigned a trip trajectory by dividing the landed weight for that species by the number of corresponding VMS records of that trip. In this process, only VMS records with speeds equal or below 3.5 knots were considered, assumed to be unequivocally associated to gear haul-up, based on existing knowledge on the operation practices in the coastal multi-gear fleet and the analysis of the frequency distribution of speed records.

Global LPUE heatmaps were thus obtained for each species. A global map of all VMS points used (all species) was also produced to indicate the area covered in this study.

## RESULTS

The three most important species for the fleet analysed and the period considered, both in quantity and landed value are the common octopus, the black scabbardfish and the European hake, representing 50% of the landings. However, while the two former species are almost exclusively landed by the multi-gear fleet, the European hake is mostly landed by coastal trawlers, the landings for multi-gear fleet representing only 40% of the landed weight for this species, and furthermore is subject to formal assessment. Therefore, it will not be analysed here. The next group of species, by order of importance of the landings in weight, are Atlantic horse mackerel, pouting, surf clam, European conger, swordfish, thornback ray and blue shark, altogether comprising around 27% of the total landings. In value, again by order of importance, the species ranking four to ten and representing 25% of the revenue are swordfish, common sole, surf clam, John dory, pouting, angler and European conger. Data with total landings in weight and value, ranks and relative importance are presented as supplementary material (Table 2A).

The five species with higher median LPUE (more than 20 kg LoA^-1^ trip^-1^) are, by decreasing order, black scabbardfish, surf clam, swordfish, Atlantic pomfret and smooth clam. Data with indicator of the distribution are presented as supplementary material (Table 3A).

The spatial distribution of the fishing activity for vessels where VMS data was available is represented in Figure 1 (the blue area is obtained by marking the VMS points identified as fishing activity).

**Figure 1.**
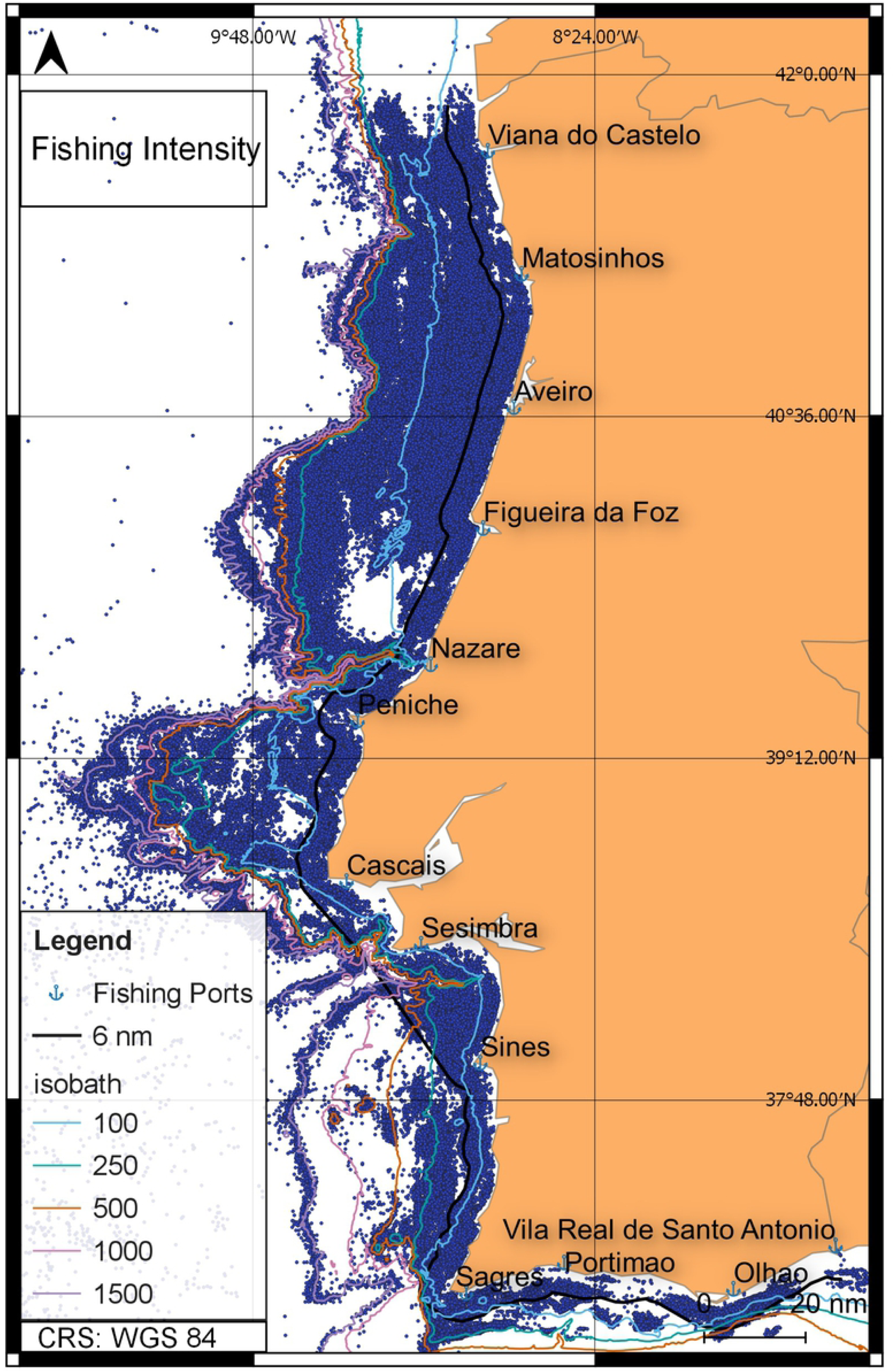
Spatial distribution of the fishing intensity of the multi-gear coastal fleet from 2012 to 2016 in mainland Portugal. Main fishing ports are indicated as well as the 6 nautical miles line (inner limit of trawling activity). Plotted pings of speeds below 3,5 kt, associated with fishing activity.

The fishing activity extends along the entire coast until the 500 metres isobath, with particular intensity to the north of the Nazaré canyon, where the continental shelf is more extended. Between Nazaré and Setúbal canyons, the bottom is characterized by numerous physiographic features (the Central Portuguese submarine canyons, Nazaré, Cascais and Setúbal – Lisbon canyons, Lastras *et al*., 2009). In these regions, there are multiple rocky areas where trawls can hardly operate, opening the opportunity for fixed gears outside the 6 nm. South of Sesimbra and along the south coast, most of the multi-gear fleet activity is inside the 250 m isobath. This is related to the intense exploitation of the continental slope by trawlers targeting mostly deep-water crustaceans. All along the coast, narrow strips following the 1000 m isobath (north of Sesimbra) and the 1500 m isobath (between Sesimbra and Sagres) and along canyons, constitute the fishing grounds for longline fisheries targeting mostly blackscabbard fish.

The results of the linear model applied to the variables YEAR, NAO INDEX, SEASON and REGION are presented in Table 1, for each of the 48 species considered, including the coefficients for the continuous variables (YEAR and NAO INDEX) and the indication of the level of each factor (SEASON and REGION) with higher LPUE. Overall, the two explanatory variables that showed a higher number of significant correlations were SEASON (41 species) and REGION (40 species). Three bivalve species were only present in the southern region, making region relevant for 43 of the 48 species considered.

**Table 1.**
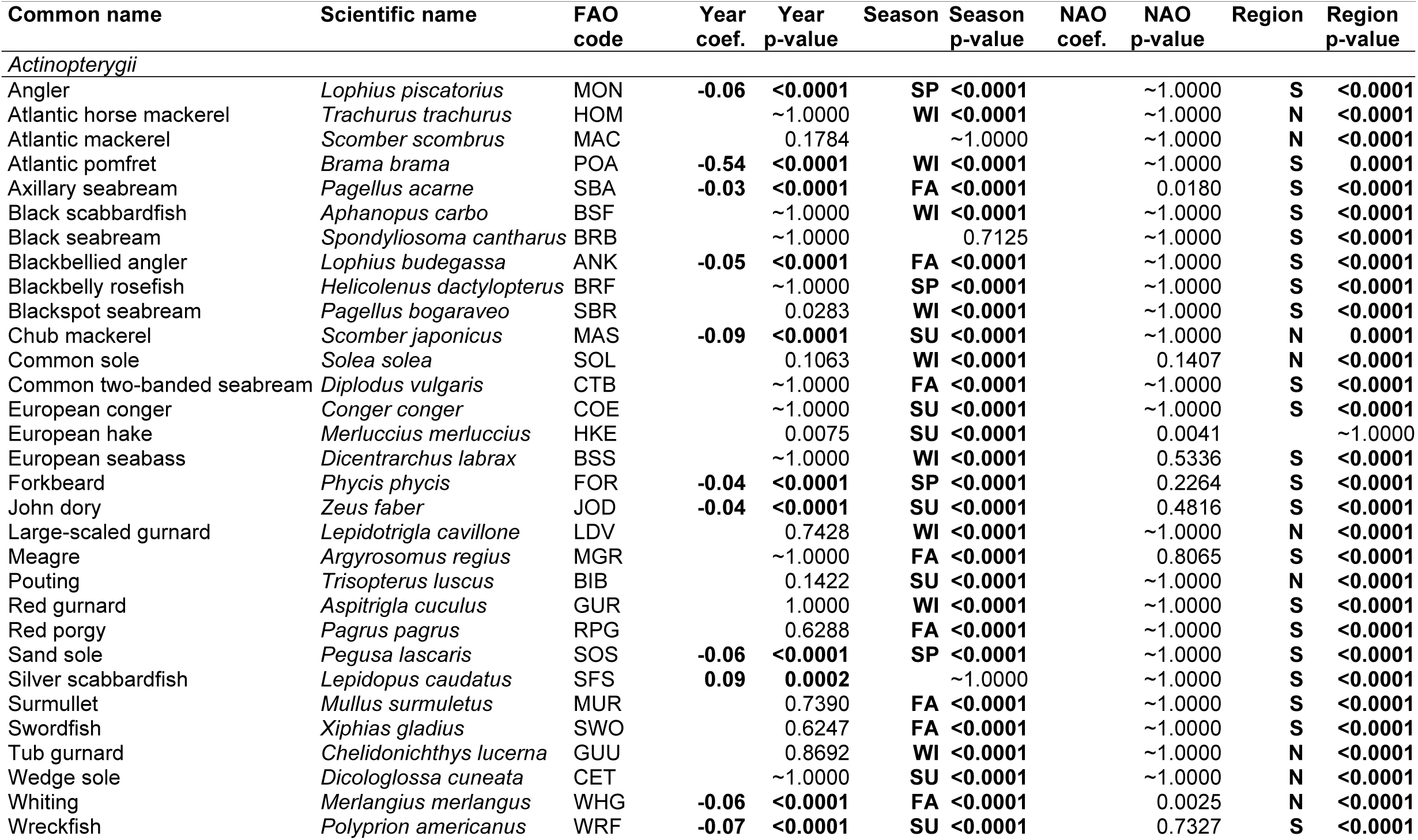
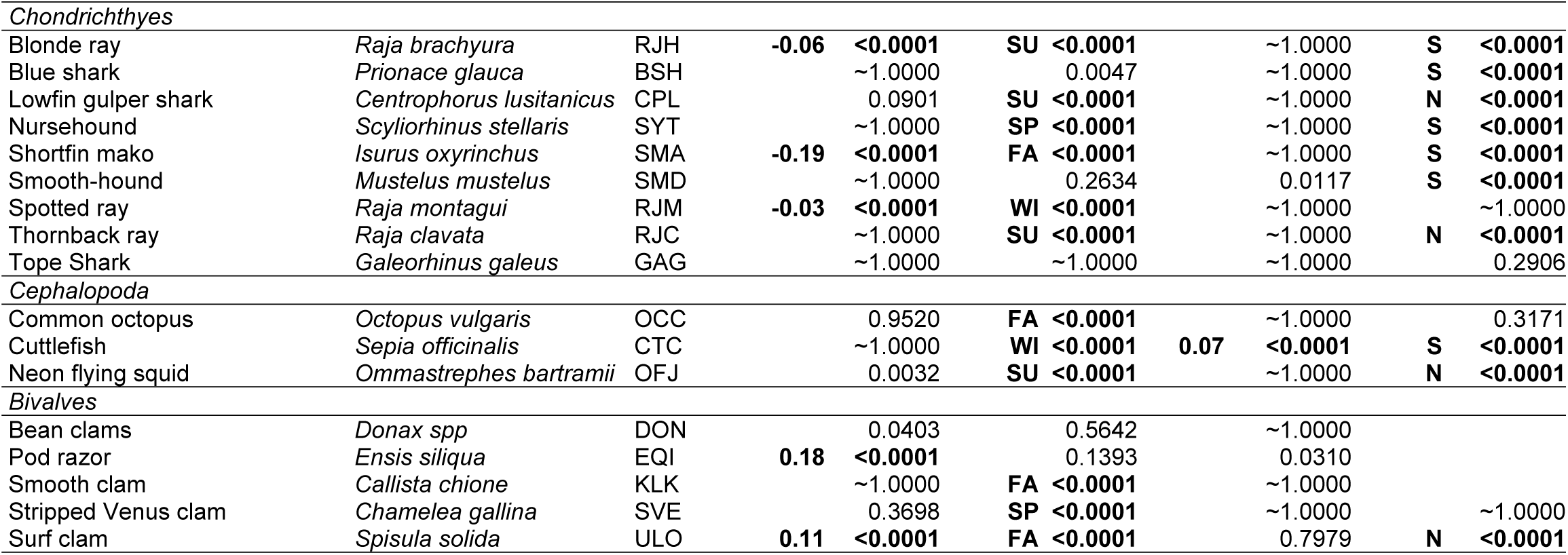
Results of the linear model for each of the 48 species considered, with year, NAO, season and region respective coefficients and Bonferroni adjusted p-values. The species are sorted by alphabetic order of their common names inside each taxonomic class. Only p-values <0.001 were considered significant. Season and region codes: FA – fall, SP – spring, WI – winter, SU – summer and N – north, S – south respectively.

A total of 18 species presented yearly trends in the period in study, of which 4 showed a positive and 14 a negative trend. NAO INDEX was significant for 3 species: European hake, whiting and cuttlefish.

A total of 31 species were analysed in the Actinopterygii class. Except for the European hake, all species presented regional differences with respect to LPUE. Nine species had significantly higher LPUE in the north (Atlantic horse mackerel, Atlantic mackerel, chub mackerel, common sole, large-scaled gurnard, pouting, tub gurnard, wedge sole and whiting), while for the remaining species LPUE was significantly higher in the south. Twelve species displayed significant interannual variations, 10 of which with negative and two with positive coefficients. The former group comprised angler, Atlantic pomfret, axillary bream, blackbelly angler, chub mackerel, forkbeard, John Dory, sand sole, whiting and wreckfish while, in the latter, silver scabbardfish were included. Seasonal variations were significant for a total of 28 species, with higher LPUEs during the winter for nine (Atlantic horse mackerel, Atlantic pomfret, black scabbardfish, blackspot seabream, common sole, European seabass, large-scaled gurnard, red gurnard and tub gurnard), during the spring for four species (angler, blackbelly rosefish, forkbeard and sand sole), during the summer for seven species (chub mackerel, European conger, European hake, John dory, pouting, wedge sole and wreckfish) and during the fall for the remaining eight species (axillary seabream, blackbellied angler, common two-banded seabream, meagre, red porgy, surmullet, swordfish and whiting). The NAO index was significant for European hake and whiting.

In the Chondrichthyes class, season was significant for seven out of the nine species analysed: blonde ray, blue shark, lowfin gulper shark, thornback ray (LPUE significantly higher in the summer), nursehound (in the spring), shortfin mako (in the fall) and spotted ray (in the winter). Geographic differences were detected for seven species, two with higher LPUE in the north, lowfin gulper and thornback ray, while the remaining five species, blonde ray, blue shark, nursehound, shortfin mako and smooth-hound, with higher values in the south. A negative trend over time was significant for three species, shortfin mako, blonde ray, and spotted ray. One species, the tope shark did not show any significant association with the explanatory variables in the model.

Season was significant for the three cephalopod species in analysis, the common octopus, the cuttlefish, and the neon flying squid, with higher LPUE in the fall, winter and summer, respectively. Regional differences were significant for cuttlefish and neon flying squid, the first more abundant in the north and the later in the south. Inter-annual variations were significant and with a negative trend for the neon flying squid. For cuttlefish, the correlation between LPUE and the NAO index was significant and positive.

Of the five bivalve species, bean, pod razor and smooth clams were exclusively landed in the south, while regional differences were detected for surf clam, with higher LPUEs in the north. Pod razor and surf clam presented an increasing interannual trend, while seasonal patterns were present for the smooth and surf clams, with higher LPUE in the fall, and for the stripped Venus clam, with higher LPUE in the spring. The bean clam did not show any significant correlation although region is implicitly important since this species was caught only in the south.

The interpretation of the significance for the important explanatory variables used in the linear model (NAO index excluded) can be visualized in Figure 2. Year and season are compounded in the variable time (represented in the x axis) and region is taken into consideration by plotting for north and south separately. From the large number of species in analysis, only eight species were selected to be examined in detail. All of these species are almost exclusively targeted by the multi-gear fleet, and none are subjected to formal assessment, being good candidates for the approach proposed in this work. Three are among the ten most important in quantity and/or value and are almost exclusively captured by the multi-gear fleet: black scabbardfish, common sole and common octopus. The blackbellied angler, forkbeard, the wreckfish and the cuttlefish are important caught species for specific segments of the multi-gear fleet. The shortfin mako is a threatened species with no formal assessment.

**Figure 2.**
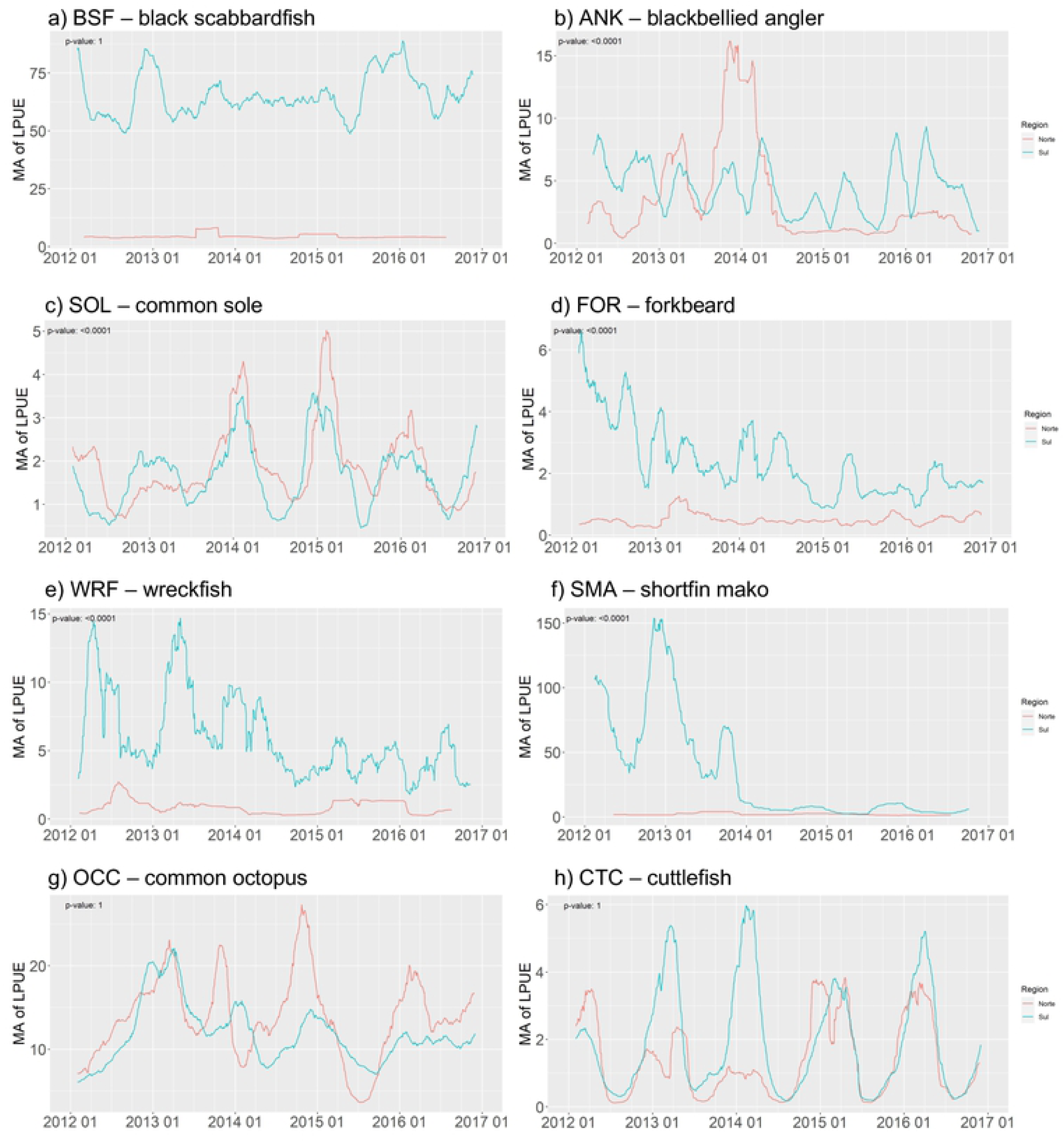
Daily landings per unit of effort (kg LoA-1 trip-1) expressed as moving average (MA) of 44 days (centre of classes plotted) for each species. The p-value of the linear trend in indicated at the top left corner of each plot. Blue and red lines correspond to the south and north regions, respectively. Species: a) black scabbardfish, b) blackbellied angler, c) common sole, d) forkbeard, e) wreckfish, f) shortfin mako, g) common octopus and h) cuttlefish.

Distribution maps of abundance (indicated by LPUE), based on landings and VMS fishing records, were produced for four of the previous species: blackbellied angler, forkbeard, shortfin mako and wreckfish (Figure 3).

**Figure 3.**
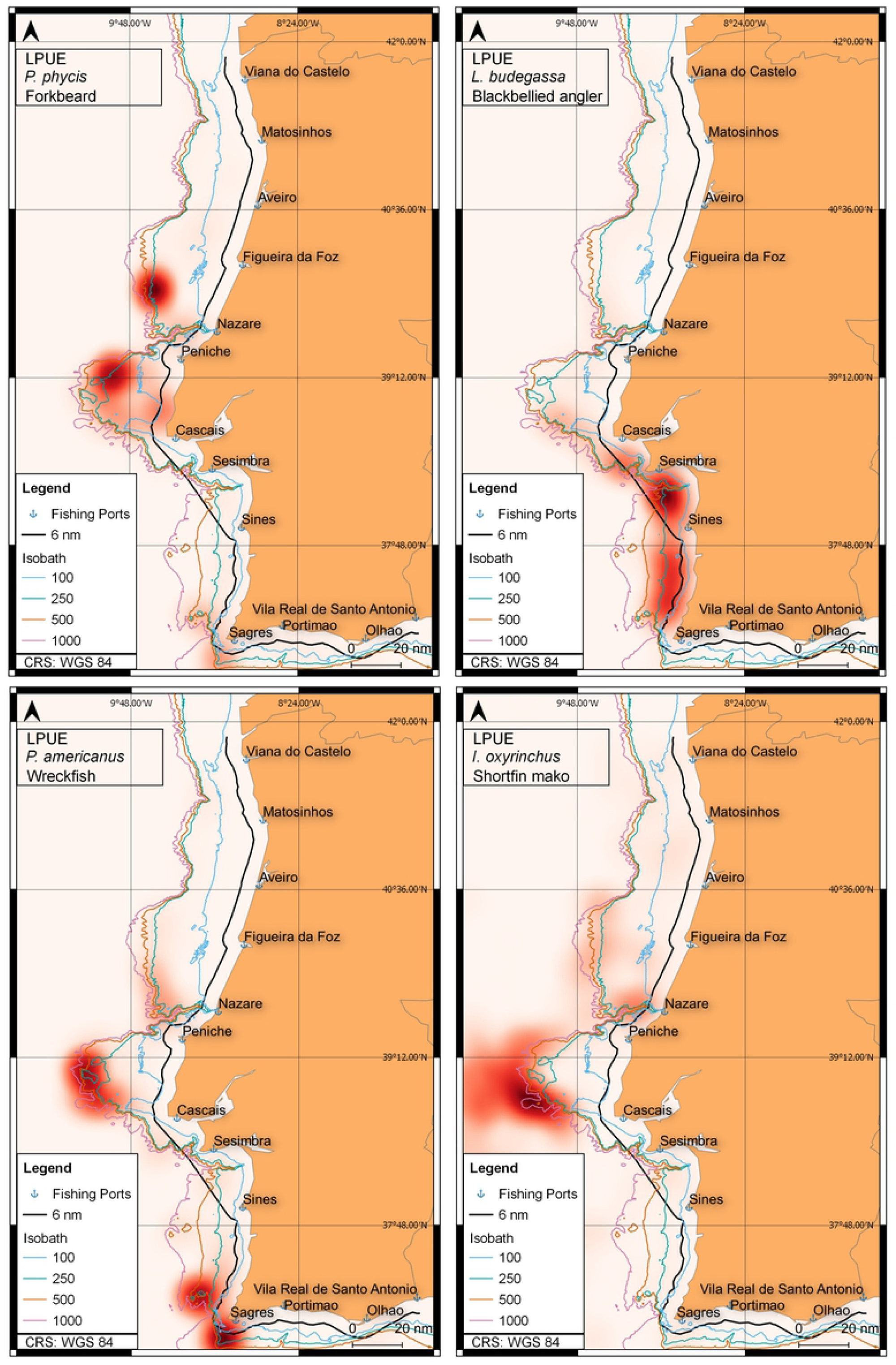
Heatmaps of abundance, using LPUE (kg LoA-1 trip-1) as a proxy, from 2012 to 2016 for a total of 4 species: a) forkbeard, b) blackbellied angler, c) wreckfish and d) shortfin mako. Main fishing ports are indicated as well as the 6 nautical miles line (inner limit of trawling activity).

The black scabbardfish is the second most important species landed. It is exclusively landed in the port of Sesimbra, being associated to the south area and despite significant higher average CPUE in the winter, it does not show clear seasonal fluctuations (Figure 2a). The blackbellied angler, with a significant reduction of LPUE over time, shows different trends in the north and south regions. The negative trend can be attributed to a significant drop in LPUE in the north after 2014 but, in the south, LPUE stayed stable (Figure 2b). For the south region, where seasonal fluctuations are clearer, there are two peaks in the spring and fall, this last one with higher average LPUE. The fishing effort for the blackbellied angler concentrates off Cascais and Pencihe while higher LPUEs are located in the Sesimbra-Sagres coast, at fishing depths between 100 and 250 meters (Figure 3a). The common sole, with significant higher LPUE in the north, displays very similar patterns with respect to season in both areas (Figure 2c), likely due to migration patterns of the species. The forkbeard, landed mainly in the south, shows a significant decreasing trend with time as evidenced in Figure 3d. Superimposed on this trend, seasonal variations can be observed, with hight average values in the spring but with peaks in early summer during most of the period in analysis. The LPUE suggests two hotspots, one North of the Nazaré canyon and a second between Cascais and Peniche (Figure 3b). Both areas are at 250m depth, close to the continental slope. The wreckfish also shows a decaying trend over time with a superimposed seasonal cycle, with higher LPUE in the summer (Figure 2e). It is also a species mostly captured in the south and is mainly concentrated in three hotspots, two off Sagres at around 250-500m depth and the third in the central region in the upper slope at depths around 500m depth (Figure 3c). A less dense area can be observed in the canyon off Nazaré.

The shortfin mako displays a distribution showing a very marked reduction 2014 onwards (Figure 2f). This protected species cannot be targeted, and the reduction may be the result of a landing ban, not necessarily a reduction in its capture. It is distributed in the central continental slope.

The two cephalopods, common octopus and cuttlefish, display typical seasonal variations related with their yearly lifecycles, with no yearly trends and higher catches in the fall and winter respectively. The cuttlefish has higher LPUE in the south, although in the last two years the values were similar in both regions, and it showed a significant correlation with the NAO index.

## DISCUSSION

The assessment and management of the species caught by the multi-gear fleet analysed here is difficult due to the large number of species (near three hundred). A simple methodology that allows the temporal and special monitoring of these resources and provides information that can be used in management is very useful, as observed in other studies with similar objectives (37).

The index capture per unit of effort (CPUE) was one of the first indicators of abundance (biomass) of a resource used in stock assessment (38), under the assumption that CPUE is linearly correlated with stock biomass and may therefore be used as its indicator. The most common phenomena affecting the proportionality between CPUE and biomass, that may invalidate its use as an indicator of biomass is hyperstability (39), which is likely to occur when fishing in areas of high concentration of the species, for example spawning aggregations (40). This situation may be combined with searching strategies targeting different concentrations of the species (41), causing CPUE to stay stable even when the resources decline. Other causes include abandonment of the fishery by less skilled fisherman when the stock declines (42) or increase in catchability due to gear improvement of better detection methods (43). The stability of the fleet in the present study, with a tendency to maintain the number of vessels and the gears used. along with the short period analyzed, contributes to reducing the probability of the referred factors to play a major role in changes in the CPUE – abundance relationship.

Accurate estimation of the fishing effort was not possible, since finer measures of effort based on the gear characteristics (i.e., net length, number of hooks or traps or soaking time) were not available. Thus, vessel length was used to standardize fishing effort. Vessel length is an indicator that reflects the fishing capacity since there is an association between vessel’s length and the maximum fleet size (number of net panels or hooks) it is allowed to operate.

In this study CPUE was replaced by LPUE, assuming that catch and landings do not differ substantially due to low levels of discards. Of the different gears used by the multi-gear fleet the most important are longlines, tangling and trammel nets, and pots. Studies in discards are available for the Portuguese continental coast for trammel nets and pelagic longlines. For trammel nets, (44–47) indicate chub mackerel (*Scomber japonicus*), sardine (*Sardina pilchardus*) and longspine snipefish (*Macroramphosus scolopax*) as the most discarded species, other species occasionally discarded being longfin gurnard (*Chelidonichthys obscura*), dragonet (*Callioynimus lyra*) and bull ray (*Pteromylaeus bovines*). For the pelagic longline (Coelho et al. 2005) the discarded species were smooth lanternshark (*Etmopterus pusillus*), lesser-spotted dogfish (*Scyliorhinus canicula*), rays (*Raja spp*) and rabbit fish (*Chimaera monstrosa*). In any case, discards were almost entirely reported for species with no commercial value. Only two of these belong to the 48 selected species in this work, the chub mackerel and unspecified rays, invalidating the use of LPUE as an index of abundance in these cases. For all the other species, the assumption can be made that LPUE can replace CPUE as an indicator of abundance.

One last important issue related to the use of global indicators of abundance such as CPUE or LPUE is the assumption that catches are a random sample of the total range of distribution of the species, meaning no selection of specific fishing grounds or areas with high density (48). This is not the case in this study, but the combination of LPUE with location (based on VMS data) can overcome this limitation allowing a spatial interpretation of CPUE.

The reason for black scabbardfish, surf clam, swordfish, Atlantic pomfret and smooth clam having higher median values of LPUE (53.5, 27.5, 21.9, 21.2 and 20.2 kg LoA^-1^ trip^-1^ respectively) can be explained due to the characteristics of the fishing and the way the LPUE index was built. Besides fishing operations resulting in high catches, LPUE are favoured by longer trips or smaller vessels. The black scabbardfish (with median LPUE above 50) is targeted by a highly efficient deep bottom longline fishery with daily trips. The swordfish is caught by drifting longlines in trips that last for several days (typically three weeks; Campos *et al*., 2019). The Atlantic pomfret was reported as being the by-catch of a longline fishery targeting hake (50) but no updated information on the possibility of this species being targeted is available, justifying specific studies. The two bivalve species have high median LPUE because they have individual daily quotas (in the order of hundreds of Kg) and the vessels have low LoA. The LPUE as expressed in this work is not a good indicator of the abundance of bivalve species due to the individual daily quota system.

The results of the linear models for LPUE as a function of year, NAO INDEX, REGION and SEASON, applied to the 48 most important species, showed a wide range of significant factors, indicating that a management strategy common to the whole fleet would be very difficult to apply. One clear output is the need to consider region when implementing management actions. In this study two regions were considered, north and south, separated by the Nazaré canyon on the west coast. While ubiquitous specie such as the common octopus showed an even distribution throughout both regions, most species were strongly associated with the north or south areas, with southern species (lowfin gulper shark, shortfin mako, black scabbardfish, forkbeard, silver scabbardfish, swordfish and wreckfish) opposed to Northern ones (surf clam and neon flying squid).

Regarding the 41 species for which significant seasonal variation was estimated, all eight selected species for further graphical analysis showed higher LPUEs towards specific seasons. In some cases, abundance patterns of one species may be related to another species, as it is the case of the common sole and the cuttlefish which are mostly caught together. While the cuttlefish spawns in inshore waters and migrates to offshore waters (∼6 nm) during autumn and winter for growing purposes (51), the common sole spawns and grows during 2 years in on-shore nurseries until moving to deeper waters from summer to autumn (52). Since the fleet’s fishing grounds for these species concentrate around the 200 meters depth, which is the maximum depth distribution of both species, our findings suggest that higher availability to fisheries during the winter periods cause higher LPUE. The blackbellied angler also shows a distinctive seasonal pattern in LPUE with a steep decrease in the autumn period and further decrease in winter. This is attributed to the existence of a fish closure between January and February, comprising the reproduction period, with the purpose to control the stock decline of anglers (*Lophius* spp.). For the Atlantic wreckfish and the shortfin mako LPUE was found to be significantly higher during the summer and fall respectively. Although caught in different seasons and with different fishing gears, both species are mainly targeted by the same, highly specific, group of vessels operating in the NE Atlantic, that often use seamounts as fishing grounds. While the bottom longline is used during the summer periods to target wreckfish over the seamounts, the use of drifting longlines is higher during fall to mainly target swordfish, capturing also pelagic sharks such as shortfin mako. (53). Since wreckfish is a sedentary territorial species and shortfin mako a large pelagic migrator it is here suggested that although wreckfish and shortfin mako are caught all year, a highly specific fleet concentrate the effort during the summer and fall seasons. Due to this, when managing the wreckfish fisheries, the difference between the coastal and seamounts fleets should be considered. The black scabbardfish shows higher LPUE levels on the winter periods. Although little is known about the life cycle of this deep water species, it is suggested that they undergo migration from the West British Isles growing grounds to lower latitudes for reproduction purposes (54). The higher LPUE associated to winter, which was also suggested in previous studies about this fleet, is yet to be explained (55). Finally, the forkbeard is mainly captured by the bottom longline fleet that targets more commercially exploited species such as blackbelly rosefish, blackspot seabream, red porgy and silver scabbardfish. Although little is known about its life cycle, the slower growth rates match higher LPUE during spring (56). At first, this could indicate that species undergo migration away from feeding grounds and increase its availability to fisheries, nevertheless it’s seasonal variability should be carefully considered due to not being a main target species.

Significant trends over time were observed for 18 species, from which four displayed an increasing trend and the remaining a negative trend during the period in study. Although statistically significant, most of these trends have little practical impact since the absolute value of the slope, for log(LPUE) as a function of year, is close to zero (for 14 species less than 0.1). Such a result is expected given the short period considered. For some species, landings were found to be restricted to some years, as it is the case of the sharks (lowfin gulper shark and shortfin mako), landed mostly from 2012 to 2014, with minor landings in subsequent years. For other species, such as neon flying squid and silver scabbardfish, peaks were registered in LPUE in 2016 and 2014-2015, respectively.

Only three species showed a significant correlation between NAO index and species relative abundance, cuttlefish, whiting and European hake. European hake is not considered in the context of this work because it is mainly landed by coastal trawlers. With respect to cuttlefish, a non-significant relationship between the NAO index and the abundance of this species was reported for the Mediterranean Sea (57) but it is know that mollusc growth is influenced by temperature (58), and a positive correlation would be expected given the positive association between higher NAO indices and sea surface temperature (59). A long-term study in the North Sea found no significant correlation between the species abundance and the NAO index (60) but in other studies, positive correlations were found. A correlation between the NAO index (warmer and dryer than average summer conditions) and the mean length of the age-0 group in the following winter was found for the population using the Bristol Channel as a nursery ground (61). A positive corelation with abundance was also found for the population of the Thames estuary (62). Better information of the spatial distribution of nursery grounds would be necessary to understand the possible relationship of the NAO index and whiting abundance off the Portuguese coast.

Species such as the common octopus, anglers, black scabbardfish, sole, swordfish and wreckfish are known to be targeted by this multi-gear fleet. Nevertheless, several species addressed in this study are being captured as by-catch. It is the case of the gurnards (tub gurnard *C. lucerna, r*ed gurnard *A. cuculus and l*arge-scaled gurnard *L. cavillone*), wedge sole, *D. cuneata* and the Atlantic mackerel *S. japonicus*, captured in small nets with small mesh size and skates (spotted ray, thornback ray, and blonde ray), caught in large mesh size nets.

Nine of the 48 most important species are Chondrichthyes species, known for their late-maturity and high longevity associated with high commercial interest which makes them very susceptible to overfishing (63), and two of them, the thornback ray and the blue shark rank 9 and 10 in importance due to amounts landed. During the period in study shortfin mako displayed a sharp decrease but this may be due to discarding in recent years. According to ICCAT, this species was reported threatened in the North Atlantic during the period covered in this work due to bycatch of the swordfish fishing fleet (64). Since 2019, the shortfin mako was introduced in the CITES appendix II and in 2020 landings were forbidden in multiple European countries due to decreasing population.

Forkbeard, which is mainly caught as bycatch by the longline coastal fleet operating in the southern coast, showed a steep decrease. This is not the case in the north, where landings are lower and constant throughout this period. A population decline was previously considered in 2011-2012 when Atlantic and Mediterranean catches dropped by 50% (65). The hypothesis of a stock decline starting in 2012 and continuing onward until, at least, 2016, the upper limit of the time range in our data, is plausible. Lastly, wreckfish, which is mainly caught by large longline vessels that operate in the Atlantic seamounts, also showed a decreasing trend during the 6-year period. This fleet mainly shifts between drifting longline, for swordfish, and bottom longline, for demersal species such as wreckfish (53). Current information regarding population trends in abundance indicates a decrease since 2015, particularly in the Azores area (66). Since the wreckfish is a long-living, late maturity species not subject to conservation measures such as minimum conservation reference size (MCRS), quotas and fishing gear restrictions or regular stock assessment, it is particularly vulnerable to over-exploitation. We hereby suggest that a monitoring program to evaluate their weight and length distributions should be carried out.

Although modelling and mapping data are from two different sources, landings and VMS respectively, both are in line with the existence of regional differences in species assemblages. A previous study (25) demonstrated the existence of two fish assemblages separated by the Nazaré canyon, which is in accordance with the results obtained for the majority of the species in our study. Regional differences, evidenced when mapping fishing effort and LPUE, can be used as indicator of changes in the fishing activity or condition of the stocks in a timely manner.

Trends in LPUE are a simple approach to characterize fisheries status in multi-species fisheries, where the high number of species involved makes the cost of fisheries independent stock assessment for all exploited species unrealistic. Due to this, when relative abundance through landings or catch data is the only data available, as it is the case for wreckfish, forkbeard, john dory and shortfin mako in this study, these indexes provide the best available information for the implementation of management advice. The combination of LPUE indexes with VMS data, can be used to monitor populations for species where discards are negligible and quota systems do not generate constant LPUE values.

The approach used here needs to be expanded by improving the available information namely increasing the length of the time series analysed (to distinguish between random fluctuations and temporal trends) and considering information on migratory and lifecycle patterns (to correctly interpretate the geographical and seasonal changes in LPUE). In this work a simplistic approach with respect to region was considered, with only two zones included in the model. Variations in fleet characteristics and oceanographic conditions, such as depth, may suggest a different partition of the area studied. The influence of the gear type on LPUE and the associations of species caught by each gear also need to be considered, maybe complemented with interviews to the skippers to identify target and by-catch species.

In conclusion, it is suggested that the methodology proposed here, consisting in the analysis of LPUE in specific areas over time, as an indicator of the fish condition, will be useful to monitor fished populations and to provide guidance for management when fisheries independent data are not available.

## ACKNOWLEDGEMENTS

This study was carried out at the scope of the project TECPESCAS – Project Mar 2020 16-01-04-FMP-0010–IPMA. The fisheries data was supplied to IPMA by the Portuguese Directorate-General for Natural Resources, Safety and Maritime Services (DGRM).

